# The impact of crystallographic plane orientation as an unexplored terrain in hemocompatible material design

**DOI:** 10.64898/2026.01.30.702901

**Authors:** Z. V. Parlak, N. Labude-Weber, A. Krause, K. Neuhaus, C. Schmidt, L. Mueller, C. Radermacher, S. Ruetten, A. Henss, S. Ferraris, S. Spriano, S. Neuss, J. Gonzalez-Julian, K. Schickle

**Affiliations:** Department of Ceramics, Institute of Mineral Engineering, RWTH Aachen University, Forckenbeckstr. 33, 52074 Aachen, Germany; Institute of Pathology, RWTH Aachen University Hospital, Pauwelsstr. 30, 52074 Aachen, Germany; Institute for Physical Chemistry, Justus Liebig University Giessen, Giessen 35392, Germany; Forschungszentrum Jülich GmbH, Institute of Energy and Climate Research, IEK-12, Helmholtz-Institute Münster: Ionics in Energy Storage, Corrensstr. 46, 48149 Münster, Germany; Politecnico di Torino, Corso Duca degli Abruzzi, 24-10129 Turin, Italy; Helmholtz Institute for Biomedical Engineering, Biointerface Group, RWTH Aachen University Hospital, Pauwelsstr. 30, 52074 Aachen, Germany

**Keywords:** Crystallographic plane, bioceramics, hemocompatibility, platelet activation, endothelialization

## Abstract

Thrombogenicity causes significant complications in the application of blood-contacting implants, requiring strategies to prevent adverse coagulation reactions. The thrombotic responses to the foreign surfaces are mainly driven by surficial factors such as surface energy, topography, and electrochemical interactions. Although anticoagulation therapies reduce the risks of clotting, patients might still encounter bleeding complications. Therefore, rather than high-risk anticoagulation therapies to counteract coagulation, it is essential to ensure hemocompatibility through the material’s intrinsic properties. Endothelialization is crucial in preventing thrombotic complications, with various strategies explored for facilitating endothelial cell adhesion and proliferation. We investigated the impact of crystallographic anisotropy on endothelial and blood cell interactions on four main planes (A-, C-, M-, and R-planes) of single crystalline alumina (*α*-Al_2_O_3_, sapphire). Employing advanced surface characterization techniques, including SIMS, KPFM and Zeta potential measurements, our study sheds light on the hemocompatibility of biomaterials considering anisotropic effects. We elucidated that the A-plane of alumina promotes endothelialization and suppresses platelet activation in contrast to other crystallographic planes. Our investigation into cell-surface interactions provides valuable insights and contributes to the advanced biomaterial design, ultimately leading to enhanced clinical outcomes.

## Introduction

Hemocompatibility holds the utmost significance and dominates cardiovascular research. Despite the advancements in cardiovascular engineering, the clinical implementation of cardiovascular implants and devices suffers from blood compatibility. Thrombogenicity remains a major concern after the implantation of prosthetic cardiovascular devices, such as catheters, vascular grafts, heart valves, artificial hearts, and pacemakers, which are designed to be in direct and permanent contact with blood [1]. In many cases, these devices cannot fulfil the intended function properly and fail due to the undesired blood-clotting response of the material. This foreign body reaction is influenced by various surface features of the biomaterial, such as surface energy, topography, and electrostatic interactions. Although anticoagulation therapies attempt to partly ensure hemodynamic stability by cell inactivating factors, patients still face risks of fatal bleeding complications including haemorrhages and gastrointestinal bleeding [2, 3]. Consequently, numerous cardiac studies focus on enhancing the surface hemocompatibility of blood-contacting implants to minimize adverse reactions and improve the overall performance and longevity of these life-saving cardiovascular devices, with a specific emphasis on minimizing platelet activation by emulating endothelial behaviour on the surface [4].

Healthy endothelialization is a well-known factor with a crucial role in inhibiting thrombotic complications, thus promoting the efficiency and durability of implants [5–7]. The primary function of endothelial cells is to provide efficient platelet and monocyte activation inhibitors, such as thrombomodulin, tissue factor pathway inhibitor, heparan sulphate proteoglycan, prostacyclin, and nitric oxide. As a result, a healthy endothelium maintains a balanced state, preventing thrombosis and serving as an ideal non-thrombogenic surface [8, 9]. A significant reduction in the platelet activation rate on endothelial-cell-covered surfaces has been demonstrated in our previous study, emphasizing the importance of promoting endothelialization to enhance cardiovascular implant hemocompatibility [10].

Over the past years, significant efforts have been made to facilitate the formation of the endothelial layer on implant surfaces through various approaches [6, 11]. Lu *et al*. designed periodic nano-micro grooves on Ti surfaces to promote endothelial cell functions [12]. Another method to enhance endothelial cell adhesion, proliferation, and migration is surface modification with extracellular matrix (ECM) molecules such as fibronectin, collagen, or vitronectin [13]. Additionally, researchers have explored the functionalization of surfaces with hydroxyl groups [14] and strong covalently grafted RGD peptides [15] to improve endothelialization.

The ability of endothelial cells to form a layer on biomaterials is mainly accelerated via adhesive biomolecules. This process involves an interplay of material chemistry, topography, wettability, and elasticity aspects [16]. Based on our previous experiments, we hypothesize that the density of atoms on the material surface and the functional groups attached to these atoms are at the origin of the sequential mechanisms. We observed a significant reduction in platelet activation and adhesion on specific crystallographic planes of single crystal alumina. Furthermore, the proteins attached to the surface also exhibited plane dependent variations [10].

Single crystalline alumina (*α*-Al_2_O_3_, sapphire) exhibits high success levels in clinical dentistry applications enabling it to withstand higher stress load compared to its polycrystalline form [17–20]. Akagawa *et al*. have demonstrated high fibrous connective tissue compatibility of single crystalline alumina [21]. Sharova *et al*. utilized sapphire as an optical port implant for photo theranostics in deep-lying brain tumours [22]. Besides focusing on its superior biological properties, this study explored how anisotropic properties influence cell behaviours. This mostly neglected anisotropic aspect in biomedical research, in fact, mediates and leads to main surficial properties such as surface potential, surface charges, and surface energy [23]. The anisotropic behaviour of alumina is currently a focal point of the research, and its role has been reported, such as in mechanical and chemical polishing [24], optical irradiance removal [25], and magnetic responses [26]. However, it is important to note that studies exploring the biological aspects of anisotropic behaviour remain limited.

In this research, we present an in-depth investigation into the influence of crystallographic anisotropy on endothelial and blood cells, which has not been explored before. The origin of cell behaviours and orientation-related properties are unclear due to the complexity of the interactions, therefore deep surficial characterization is essential for a thorough understanding of the interface. To achieve this, we meticulously examined four main planes of single crystalline alumina, specifically 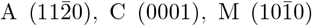, and 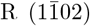 (Figure 1). This comprehensive analysis offers significant guidance for reliably determining the anisotropic nature of hemocompatible biomaterials and thus is expected to contribute to the advancement of biomaterial design. The potential implications of our research extend to the enhancement of endothelialization and the overall improved performance of biomaterials in various clinical applications.

**Figure 1.**
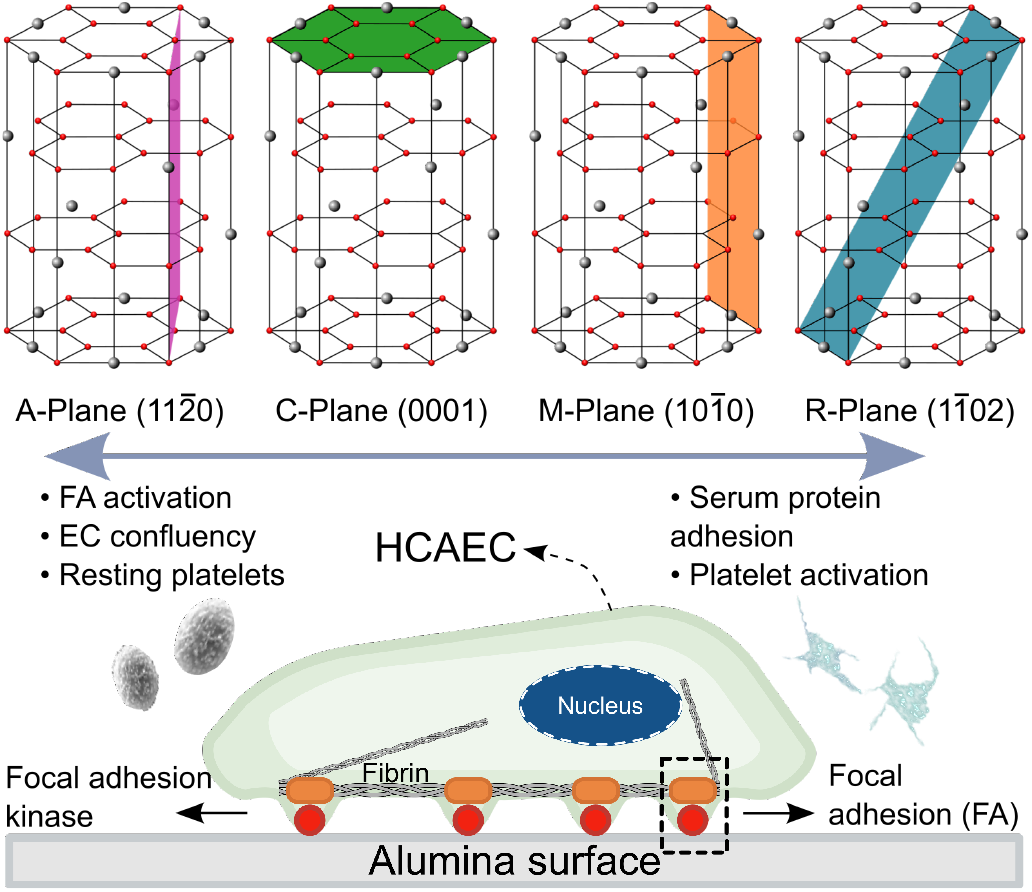
Crystal structure of hexagonal single crystalline alumina and 3D arrangement of Al^3+^ (Gray) and O^2−^ (Red) ions. Crystallographic planes are represented in different colour codes (above). Endothelial cell adhesion on the alumina surface, which is mediated by adhesion biomolecules, is illustrated below. Crystallographic plane-dependent trends of cellular behaviours are depicted by arrows.

## Experimental

### Structural and surface characterization

Aluminium oxide (*α*-Al_2_O_3_) single crystals (5 mm x 5 mm x 1 mm) were commercially provided by Mateck GmbH (Jülich, Germany). The specimens were cut in four crystallographic planes 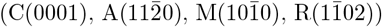 and polished on both sides. To confirm the crys-tallographic orientations, x-ray diffraction (XRD) analysis was conducted using an Empyrean™ 3rd Gen. diffractometer from Malvern PANalytical, featuring an ICore optical system and a GaliPIX3D area detector. The divergence was set to 0.25 degree, and masks were arranged to illuminate a spot of approximately 10 mm in diameter. The powders were positioned on a zero-background-holder made of a silicon single crystal. Following the analysis, the crystalline phases were verified using Highscore Plus software, and structural data were obtained from COD and ICSD databases.

Surface roughness and surface potentials were simultaneously assessed using a Cypher ES Atomic Force Microscopy (AFM) system (Oxford Instruments, UK) equipped with Pt-coated PPP-NCST-Pt tips (Nanosensors, Switzerland). The measurements were conducted under atmospheric conditions at a consistent sample temperature of 37°C, mirroring the physiological thermal conditions of the human body. The determination of surface potential was performed by Kelvin Probe Force Microscopy (KPFM) in a dual-pass experiment. In the initial pass, sample topography was mapped using intermittent contact mode, while in the second pass, the surface potential was measured along the same line, maintaining a fixed average height difference (30 nm) between the vibrating tip and the sample surface. Mean square height (Sq) and the developed interfacial area ratio (Sdr) were computed for a series of topography images under ISO 25178 standards. Measurements were taken at multiple locations on each plane to ensure statistical validity.

To assess surface wettability, dynamic contact angle measurements were executed utilizing the tangent 2-method on a drop-shape analyser system (DSA 100, Krüss, Germany). Prior to these evaluations, the specimens underwent sequential rinsing procedures involving ethanol and distilled water to mitigate potential contamination effects. A precisely controlled volume of distilled water (3 *µ*l) was dispensed onto the sample surface at a dosing rate of 250 *µ*l/min. The average contact angle values were statistically calculated over three droplets, each measured at distinct locations on three specimens for each specimen group.

Zeta potential measurements were carried out using an electrokinetic analyser (SurPASS, Anton Paar - Graz, Austria) with an adjustable gap cell. The samples were fixed in the cell by positioning the surfaces to be investigated over an area of 10 mm x 5 mm, facing each other to create a gap, with an adjustment of approximately 100 *µ*m. The electrolyte used was an aqueous solution of 0.001M KCl and the pH titration was carried out using 0.05M HCl and 0.05M NaOH. The measurement was started at the pH of the electrolyte (5.6) and the initial curve was generated by titrating the solution in the acidic range using the instrument’s automatic titration unit. After cleaning the instrument and the samples with ultrapure water, the basic curve was obtained by starting again with the electrolyte at pH 5.6 and titrating within the basic range.

Measurements were conducted on epitaxially polished samples using an M6 Hybrid instrument (IONTOF GmbH, Münster, Germany) equipped with a 30 kV bismuth (Bi) primary ion cluster gun for analysis, a dual source column, and a gas cluster source for depth profiling. Spectra were recorded in the spectrometry mode of the LMIG with Bi^3+^ ions (FWHM m/Δm = 8360 at m/z = 26.00 (CN^−^)) and the all-purpose mode of the analyser. Regions of 500×500 *µ*m^2^ were measured for each crystallographic orientation in random mode. To ensure comparability, the dose density for all scans was maintained at 5 × 10^11^ ions/cm^2^. The spectra were calibrated to carbon ions (C^−^, C^2−^, C^3−^). Before additional analysis, a gentle cleaning step was performed to remove surface contaminants such as adsorbed hydrocarbons using the GCIB. 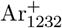 clusters with an energy of 10 kV were chosen for this cleaning process. The obtained data were evaluated using SurfaceLab 7.3 software (IONTOF GmbH). All samples were electrically isolated from the sample holder and measured with electron neutralization from the flood gun.

### Biological characterization

The cytocompatibility assessment of the single crystalline alumina was verified using Human Umbilical Vein Endothelial Cells (HUVECs) and Peripheral Blood Mononuclear Cells (PBMCs), as described in a previous publication [10]. In this study, the potential of single crystal alumina samples to promote cell viability and proliferation is tested with Human Coronary Artery Endothelial Cells (HCAECs). Before the experiments, all samples were sterilized by exposure to ultraviolet light for 15 minutes in a sterile tissue culture hood, followed by immersing in 70% ethanol for 5 minutes and finally rinsed with phosphate-buffered saline (PBS).

### Cell culture

The cryo-preserved stocks of HCAECs (PromoCell, Germany) were thawed at room temperature and cultured in EGM-2 medium containing 5% fetal bovine serum (FBS) (BulletKit, Lonza, Germany) in T75 cell culture flasks. The medium was refreshed every 3 days and cultures were maintained in a humidified incubator at 37°C, with 5% CO_2_. When the cells reached 80–90% confluence (10-14 days), the culture medium was removed, and the adherent cells were rinsed using 10 ml of PBS. Following this step, the cells were trypsinized (3 ml, 0,05% Trypsin-EDTA). After 3 minutes of incubation at 37°C, the trypsin reaction was stopped by adding 7 ml cell culture medium. Centrifugation was applied at 200g for 5 min, the supernatant was completely removed, and the cell pellet was resuspended in a defined volume of fresh culture medium. Finally, HCAECs were seeded onto sterilized samples with a density of 5×10^4^ cells/cm^2^ at passages 2 through 6. Seeded samples were analysed for the Phosphorylated Focal Adhesion Kinase (pFAK) and live cell imaging experiments described in the following sections.

According to the ethical requirements, all blood samples were collected upon the written consent of the donors. The donors received a detailed explanation of the research, they confirmed their compliance with the conditions as described in ISO 10993-4 standards. For the isolation process, peripheral blood (30 ml) was withdrawn with a Safety-Multifly needle in 7.5 ml S-Monovette Plasma Lithium-Heparin tubes and then immediately transferred into a 50 ml Falcon tube. 3 ml of blood was added into a 15 ml Falcon tube with an equal volume of Ficoll solution (Sigma-Aldrich, Germany), which is a hydrophilic polysaccharide to separate layers of blood. We note that any physical stress on the cells was strictly avoided, which might trigger cell activation. The solution was centrifuged at 400g for 30 minutes. After this centrifugation step, the blood was separated into erythrocytes, Ficoll, PBMCs, and plasma. Isolated PBMCs (including platelets and leukocytes) were transferred into a 50 ml Falcon tube and washed three times using an EGM-2 medium, by centrifuging at 250g for 10 minutes. After washing, the PBMC pellet was resuspended with an EGM-2 culture medium and seeded on the samples in 48-well plates. Approximately 10^6^ cells/cm^2^ were seeded and incubated for 4 hours.

The samples were incubated in Bovine Serum Albumin (BSA) in a concentration of 20 *µ*g/mL for 30 minutes at 37°C. Following the incubation, adsorbed protein concentrations on the sample surfaces were quantified by comparing a set of protein standards using a Micro BCA Protein Assay Kit according to the manufacturer’s instructions (Thermo Fisher Scientific). The same procedure was applied to the samples which were gently washed with PBS after incubation. The light absorbance of the protein solutions was measured by a Tecan microplate reader at 562 nm. The protein concentration values were analysed based on the standard curve plotted for each sample. A set of diluted standards as three replicates of each dilution and three samples for each plane were included in the test procedure for increasing the statistics.

HCAECs were seeded onto specimens at a density of 1 × 10^5^ cells/cm^2^ and incubated at 37°C for 60 minutes. Following the incubation period, the cells were fixed with 4% paraformaldehyde solution for 5 minutes at room temperature, and permeabilization was achieved using 1% Triton-X-100 for 5 minutes to allow for intracellular antibody penetration. Before primary antibody staining, the HCAECs were blocked with 3 wt.% BSA for one hour at room temperature to minimize non-specific antibody binding. Diluted anti-pFAK monoclonal rabbit antibody (Thermo Fisher, USA) at a ratio of 1:1000 in 3 wt.% BSA was added to the specimens and incubated overnight at 4°C to facilitate specific binding of the antibody to phosphorylated FAK proteins. Following three washes with 3% BSA in PBS to remove excess primary antibody, specimens were incubated with secondary antibody, IgG Alexa 555 (Thermo Fisher, USA), diluted at a ratio of 1:200 in 3% BSA for 60 minutes at room temperature in the absence of light. After washing to remove unbound secondary antibody, specimens were stained with Phalloidin Alexa 488 (Thermo Fisher, USA) at a ratio of 1:250 to visualize F-actin and with DAPI (Carl Roth, Germany) at a ratio of 1:1000 for nuclear staining. Specimens were stored at 4°C in dark conditions until further analysis or imaging.

Specimens were examined under a fluorescence microscope (Axio Imager, Zeiss, Germany) equipped with appropriate filter sets for DAPI (blue, excitation: 461 nm), Actin (green, excitation: 488 nm), and pFAK (red, excitation: 555 nm) channels. Two specimens per crystallographic plane were analysed using imaging software ZEISS, ZEN (Blue edition 3.4). For all analyses, at least ten cells from 3 different donors were randomly chosen and evaluated.

The viability and proliferation of HCAECs on the samples were assessed using the Cellcyte X (Cytena, Germany) live cell imaging system. Before imaging, cells were seeded onto the samples and stained with Celltracker Green CMFDA Dye (Thermo Fischer, Germany) at a ratio of 20 *µ*l per 10 ml of culture medium, followed by incubation at 37°C for 15 minutes. After staining, the alumina samples were transferred to a new well plate containing 2 ml of fresh culture medium to remove any non-adherent cells from the sample surfaces. The well plate with the cellularized materials was then placed into the Cellcyte X system. Imaging was conducted using nine adjacent, non-overlapping fields of view arranged in a 3 x 3 grid. Specifically, areas containing the cellularized material were selected for analysis. Imaging in the green fluorescent channel utilized an exposure time of 150 ms and a gain of 2 dB. Each field of view was imaged once every hour over 48 hours in both the ‘enhanced contour’ (bright-field analog) and green fluorescent channels. To maintain consistency, the cell culture medium was not exchanged during the 48-hour imaging period to avoid interrupting the process. Confluency analysis was carried out using Cellcyte Studio software, focusing on the fields of view containing the cellularized material. The surface area covered by endothelial cells on the material was quantified and exported for comparative analysis.

Philips XL 30 SEM equipped with a field emission gun, was employed to examine the cell morphology, adherence, and activation of alumina single crystals. Sample preparation for SEM involved fixing incubated cells with 3% glutaraldehyde without prior washing steps, followed by storage at 4°C. Subsequently, samples were first washed with hexamethyldisilazane and then dehydrated with 30, 50, 70, and 90% ethanol respectively. The dried samples were covered with gold-palladium coating before being scanned utilizing secondary electrons.

Platelets are anuclear blood cells, measuring approximately 2 *µ*m in size, and are typically found circulating in the bloodstream. They possess various cytoplasmic granules, some of which discharge their contents upon activation and change their morphology. They were classified based on morphological alterations, including disc-shaped, granular, spread, and fully spread forms. Granular platelets exhibited heightened contrast under scanning electron microscopy (SEM), distinguished by their granular structure positioned higher from the surface. This surface roughness resulted in a lower intensity of the e-beam, rendering them brighter [27]. Disc shapes lacking filopodia were indicative of non-activated cells. Conversely, other morphological variations suggested activation, with disc-shaped cells bearing thin, short filopodia representing an initial stage of activation for observation of activation rates (Figure 9). ImageJ software’s cell counter marker was utilized for analysis, assessing 2 samples from 3 different spots and involving 3 donors per system to ensure a robust statistical analysis. A standardized area of 85×128 *µ*m^2^ (10880 *µ*m^2^) was evaluated across all charges.

Mean experimental data with standard deviation (SD) are presented in this report. Statistical analysis was performed using one-way ANOVA coupled with Tukey’s multiple comparison test via GraphPad Prism version 9.0.0 for Windows, developed by GraphPad Software, San Diego, California, USA. A significance level of p *<*.05 was considered statistically significant.

## Results

### Intrinsic surface characterisation

Textured alumina planes (A, C, R and M) and crystal structure were confirmed by XRD before further surface characterization. All planes were identified by the main peaks observed in the XRD patterns, which precisely matched with the ICSD reference data.

Figure 2 shows AFM images of each plane. The mean surface roughness (Sq) and standard deviation of roughness values measured over the scanned area are provided. Variations in surface roughness properties among the different crystallographic planes are visible. We note that the particles adsorbed on the surfaces were excluded from the roughness analysis. The A plane exhibited the lowest mean surface roughness (1.95 nm) with a relatively low standard deviation (0.06%), indicating a relatively smoother surface. In contrast, the C plane showed the highest mean surface roughness (5.27 nm) with a higher standard deviation (0.11%), suggesting a more irregular surface morphology. The M and R planes exhibited intermediate roughness characteristics compared to the A- and C-planes.

**Figure 2.**
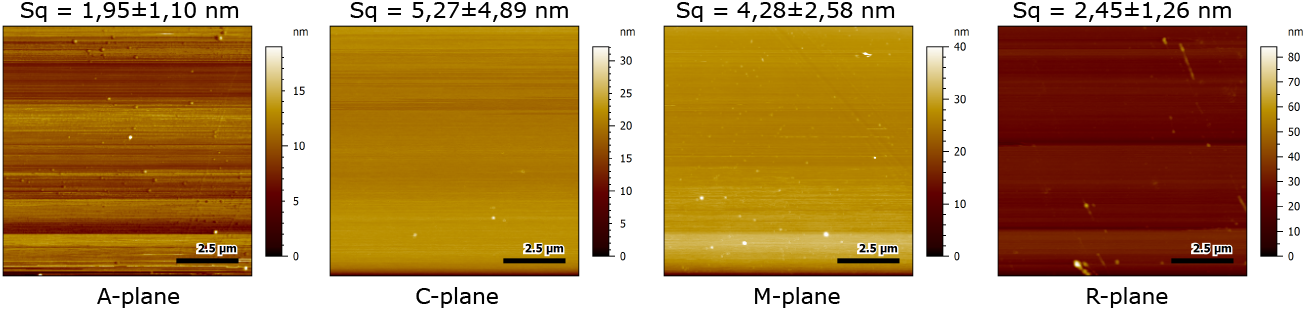
Surface topography of single crystalline samples measured by atomic force microscopy. The mean square height of an image (Sq) is calculated over an area of 100 *µ*m^2^.

The contact angle measurements revealed distinct differences in surface wettability among the different crystallographic orientations. The A-plane exhibited the highest contact angle (94^°^), indicating a hydrophobic region with low wettability. In contrast, the C-plane demonstrated the lowest contact angle, signifying high surface energy and increased wettability due to disrupted intermolecular bonds and a higher affinity for hydroxyl (OH) groups on the surface.

The surface potential measurements conducted in this study were qualitative, providing insights into the relative surface potentials of different crystallographic planes. The R-plane, with an absolute surface potential value of -114 mV, served as the reference point, assigned a surface potential of 0 mV. From this baseline, the surface potentials of other crystal planes were evaluated and compared.

Figure 4 presents the average surface potentials determined from each crystallographic plane. The A-plane exhibited the highest surface potential among the evaluated planes, with an average value of 544 ^±^ 96 mV. In contrast, the C-plane showed a significantly lower average surface potential of 110 ^±^ 35 mV. The M-plane displayed an intermediate surface potential of 122 ^±^ 30 mV. On the R-plane, stripy potential map was observed because of the strong charging leading to the repulsion of the cantilever and resulting in increased standard deviation. Regarding the surface topography, more homogenously distributed small surface impurities on C-plane are significant.

**Figure 3.**
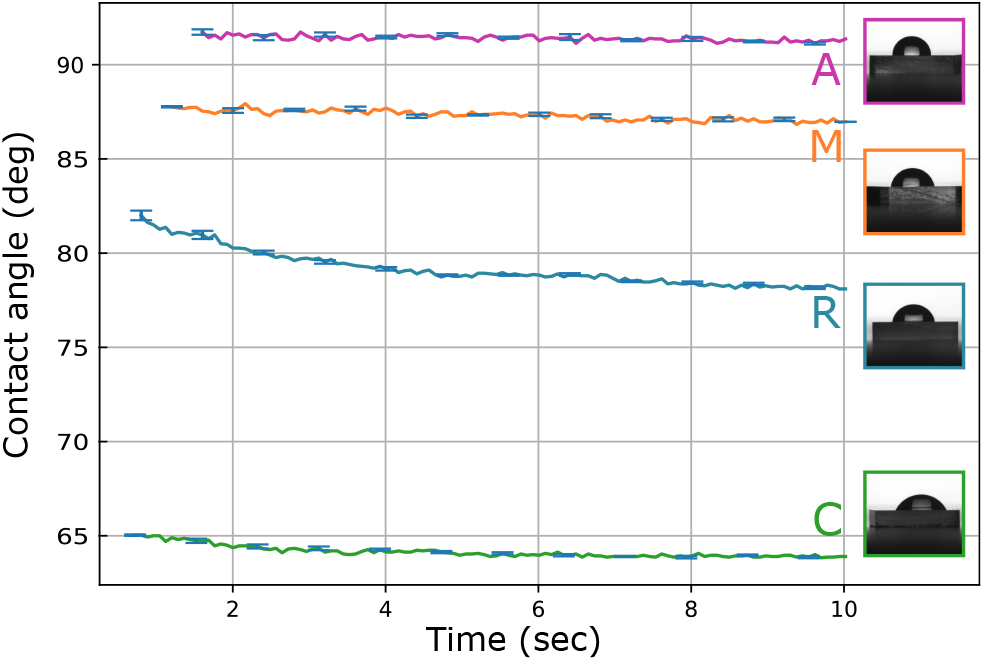
Time dependency of water contact angles and drop profiles for various planes of alumina single crystals. Error bars (standard deviations for 3x measurements and 3x regions) are given for every 10 data points for the sake of visibility.

**Figure 4.**
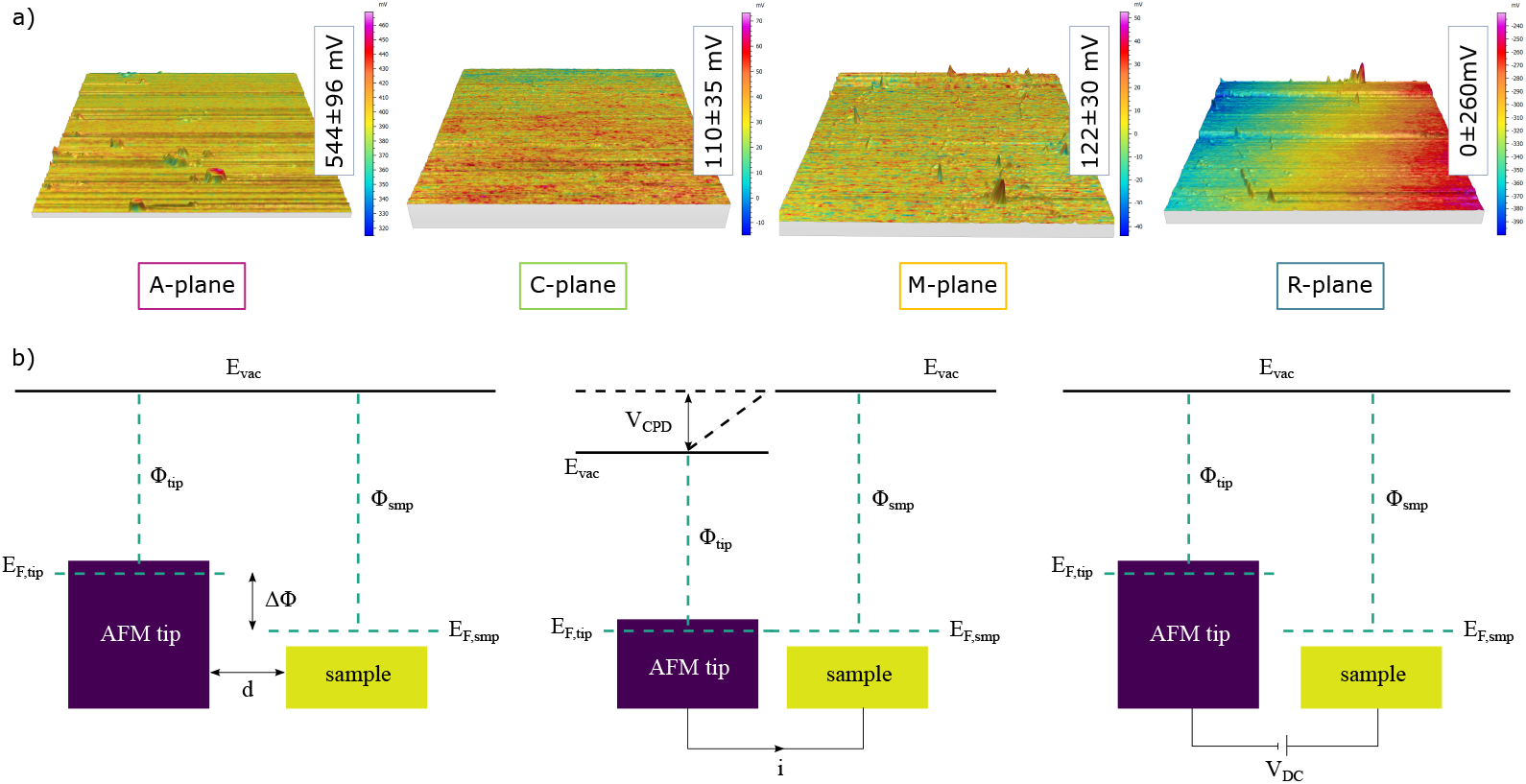
Illustration depicting surface potentials of the crystals and the AFM tip following contact and charge transfer. Below are exemplary surface potential maps obtained through KPFM measurements. The principal concept of KPFM is schematized below.

The titration curves of the A-, C-, M-, and R-planes presented in Figure 5, displayed a consistent pattern characterized by an isoelectric point (IEP) within the range of 4.1-4.6, indicative of a minimal presence of exposed functional groups with acid-base behaviour. An IEP of 4 is expected in the absence of functional groups, with almost neutral functional groups, or with a balanced amount of acidic and basic functional groups in the case of zwitterionic compounds. Although the general trends were similar among the samples, discernible differences were observed. The planes C, M, and R showed IEP around 4.1 and a plateau for pH values higher than 7.5, pointing to the presence of weakly acidic functional groups that were fully deprotonated only at high pH levels (7.5-8). The zeta potential at the physiological pH (7.4) was significantly negative (from -50 mV to -35 mV). No plateau was evident in the acid range. In contrast, the titration curve of plane A diverged from the others, with a slightly but significantly higher IEP, a shift of the onset of the plateau in the basic range to lower pH values, the presence of a small plateau in the acidic range, and a lower zeta potential value at the physiological pH (-30 mV).

**Figure 5.**
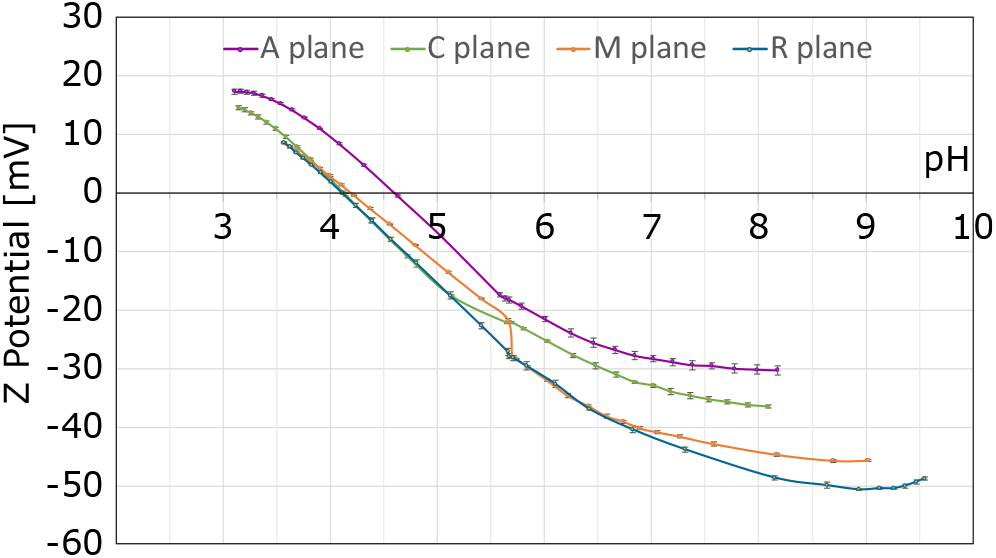
Zeta potential titration curves of crystallographic planes.

Although the general trends were similar among the samples, discernible differences were observed. Samples C, M, and R show a plateau for pH values higher than 7.5, pointing to the absence of strong acidic functional groups that would have been fully deprotonated even at lower pH levels. Instead, the presence of weak acidic groups, fully deprotonated only at higher pH (7.5-8), was inferred. In contrast, the titration curve of sample A diverged from the others, with a slightly but significantly higher IEP, a shift of the plateau onset to lower pH values in the basic range, and the presence of a small plateau in the acidic range. Five different OH groups with different chemical reactivity are identified: basic groups (pKa around 9), weakly basic (pKa at 6.6-6.8), almost neutral (pKa at 4.4-4), and acidic (pKa around 3).

For further insight into the surface chemistry, four planes were analysed using secondary ion mass spectrometry (SIMS). Due to variations in their crystallographic orientation, the stacking orders of aluminium (Al) and oxygen (O) differ, potentially influencing the levels of impurities and adsorbed molecules. Across all crystal orientations, the spectra revealed the presence of various 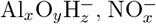, and 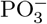 ions in the negative ion mode, alongside contaminations such as Na^+^ and K^+^ in the positive ion mode. A statistically significant trend between crystal planes was observed for 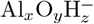 fragments. The normalized intensities of Al_2_O_3_(OH)^−^ relative to the highest intensity are shown in Figure 6. Crystal plane A featured the lowest intensity compared to all other plane orientations (p *<* 0.05). It was followed by the C-plane which displayed significantly lower intensity than M plane. The latter exhibited the highest intensity. For all four orientations the order of intensities for 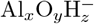 signals correlated well with the platelet behaviour, but also wettability, see Figure 3. The crystallographic effect on the hydration of the sapphire surface appeared to have a significant effect on hemocompatibility.

**Figure 6.**
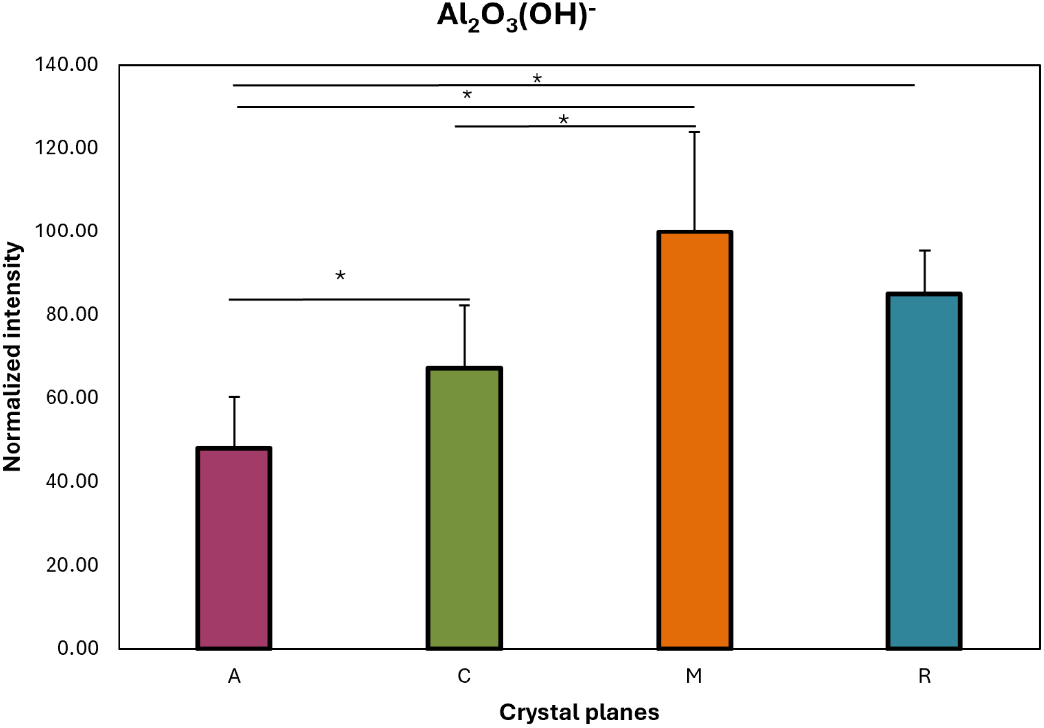
Relative abundance of the most prominent 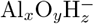 peak, Al_2_O_4_H^−^, scaled to the highest intensity of the crystal planes, M. The intensities were taken from a 50 x 50 *µ*m^2^ region within at least three different field of views and normalized to total primary ion dose. Standard deviations were used to calculate the error bars using error propagation. (*p *<* 0.05).

### Cell Behaviours

Endothelial cell proliferation was meticulously examined on different plane surfaces, revealing notable differences in adhesion intensity, and confluency. Our investigation illustrates that endothelial cells showed higher confluency on the A-plane throughout a 48 h incubation period (Figure 7). Specifically, the hierarchical order of cellular confluency among the planes was delineated as A *>* C *>* M *>* R.

**Figure 7.**
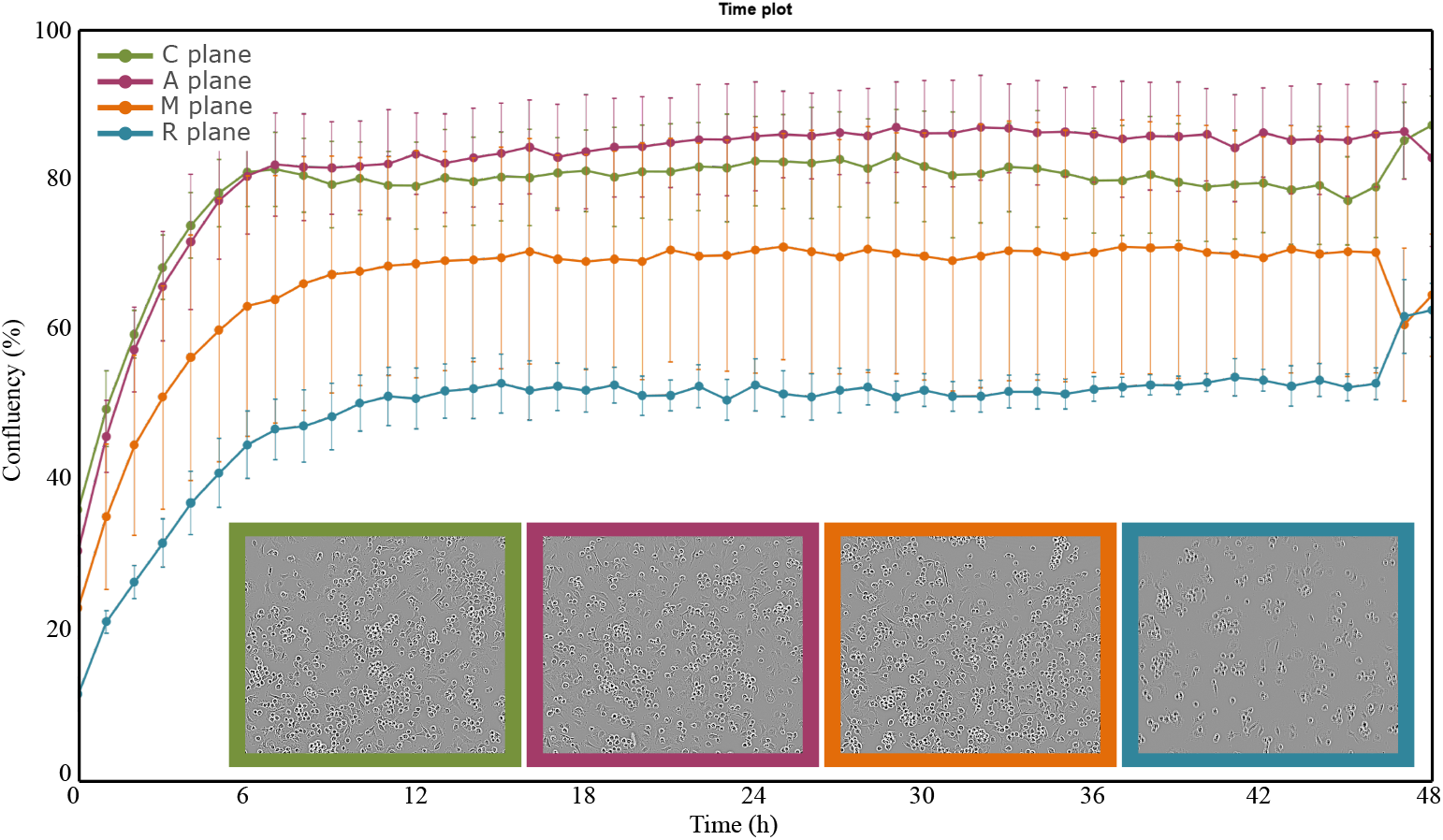
Cell confluency plots for live cell imaging over 48 h incubation period of HCAECs on different planes. The confluency data were calculated for one exemplary donor and represent 9 regions of interest over the samples. Representative images at the 12th hour of incubation for corresponding planes are given below.

The PFAK analysis images are depicted in Figure 8. Visual assessment based on the phosphorylation intensity of FAK revealed that HCAECs on Plane A exhibited a density of adhesion points. Focal adhesions were observed along the cellular membrane for 60 min after seeding. Analysis indicated that red excitation was prominently detected on Plane A, consistent with the cell confluency findings. In contrast, R-plane displayed very weak pFAK expression, signifying weaker HCAECs adhesion on the surface.

**Figure 8.**
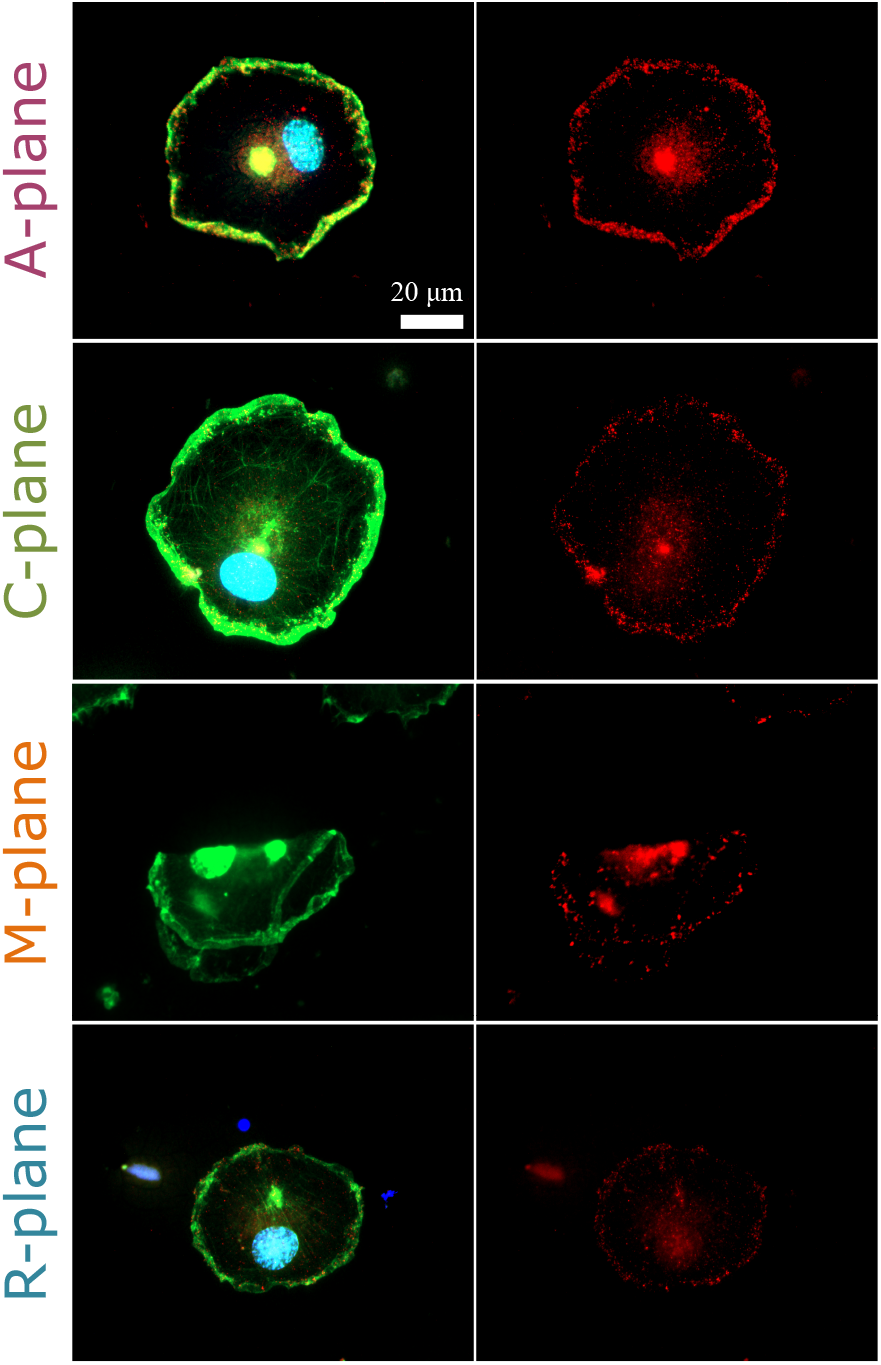
Cell confluency plots for live cell imaging over 48 h incubation period of HCAECs on different planes. The confluency data were calculated for one exemplary donor and represent 9 regions of interest over the samples. Representative images at 12th hour of incubation for corresponding planes are given below.

Platelet behaviours were examined to assess their degree of activation through SEM analysis. Figure 9 (Left) illustrates the activation states of platelets on different crystallographic planes. The analysis revealed significantly lower activation levels of platelets on the A-plane surface compared to the other surfaces. This observation suggests a discrepancy in hemocompatibility, with the M-plane surface demonstrating a higher number of activated platelets. In Figure 9 (right), a statistical analysis of the activation ratio is presented. The activation ratio was calculated as the ratio of activated cells to the total number of adherent cells. This analysis unveiled a significant influence of the crystallographic plane on the context of coagulation, as evidenced by the variation in activation ratios across different planes.

**Figure 9.**
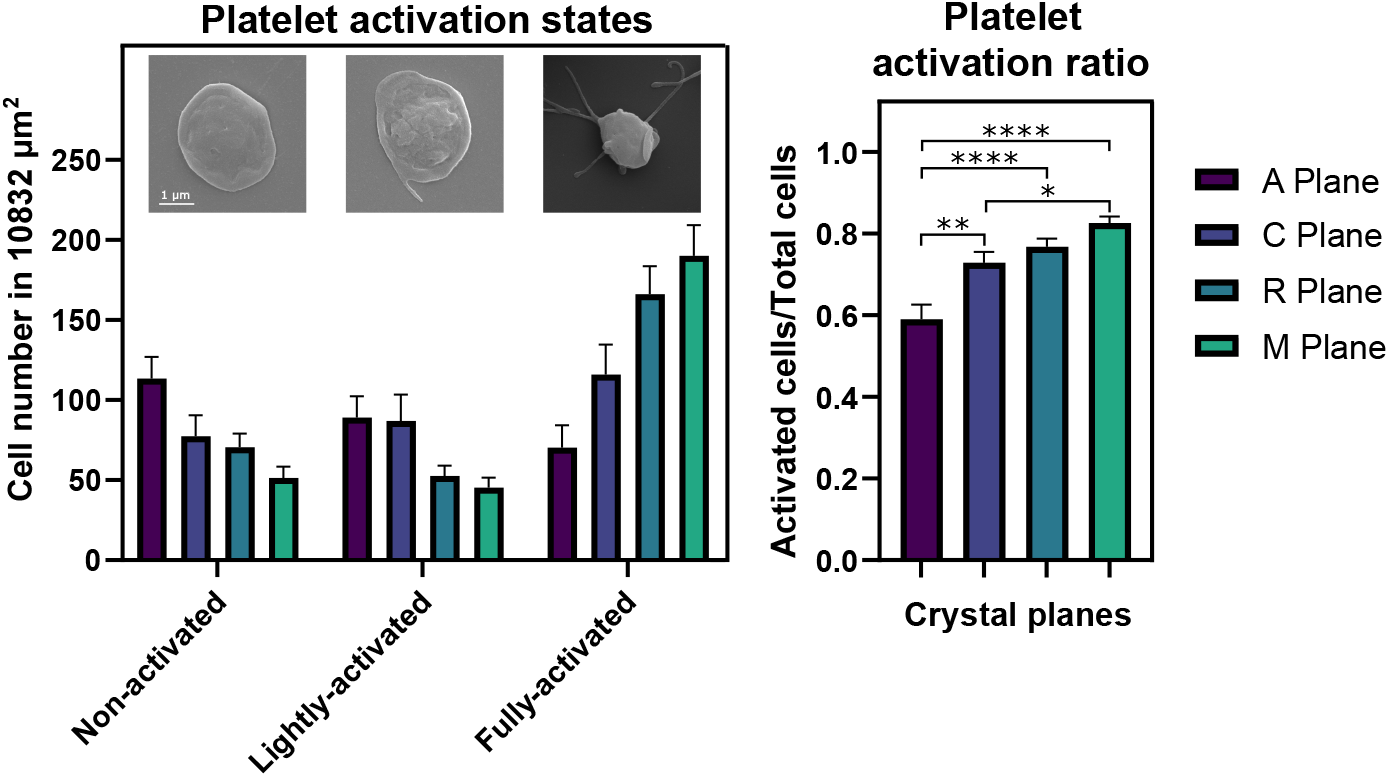
(Left) The graph indicates the activation states of platelets after 4 h incubation on alumina planes under static conditions. Each within an 85×128*µ*m2 area of interest. SEM images of cell morphology changes during activation are given above. (Right) The graph indicates platelet activation ratios. Standard deviations on the graph are calculated from one representative experiment over 18 SEM images per system, *p *<* 0.05, **p = 0.001, ****p *<* 0.0001).

Protein adhesion was quantified using the Micro BCA Protein Assay to evaluate the adsorption profile of serum proteins on the sample surfaces. Figure 10 presents the adsorption profiles of serum proteins before and after washing from the surfaces. Following the washing process, the average concentrations of protein adsorption were determined to be 9.57 *µ*g/cm^2^ on the A-plane, 9.61 *µ*g/cm^2^ on the C-plane, 11.77 *µ*g/cm^2^ on the R-plane, and 15.06 *µ*g/cm^2^ on the M-plane. Although the differences were minor, it is noteworthy that the A-plane exhibited the lowest protein adsorption, followed by the C-plane.

**Figure 10.**
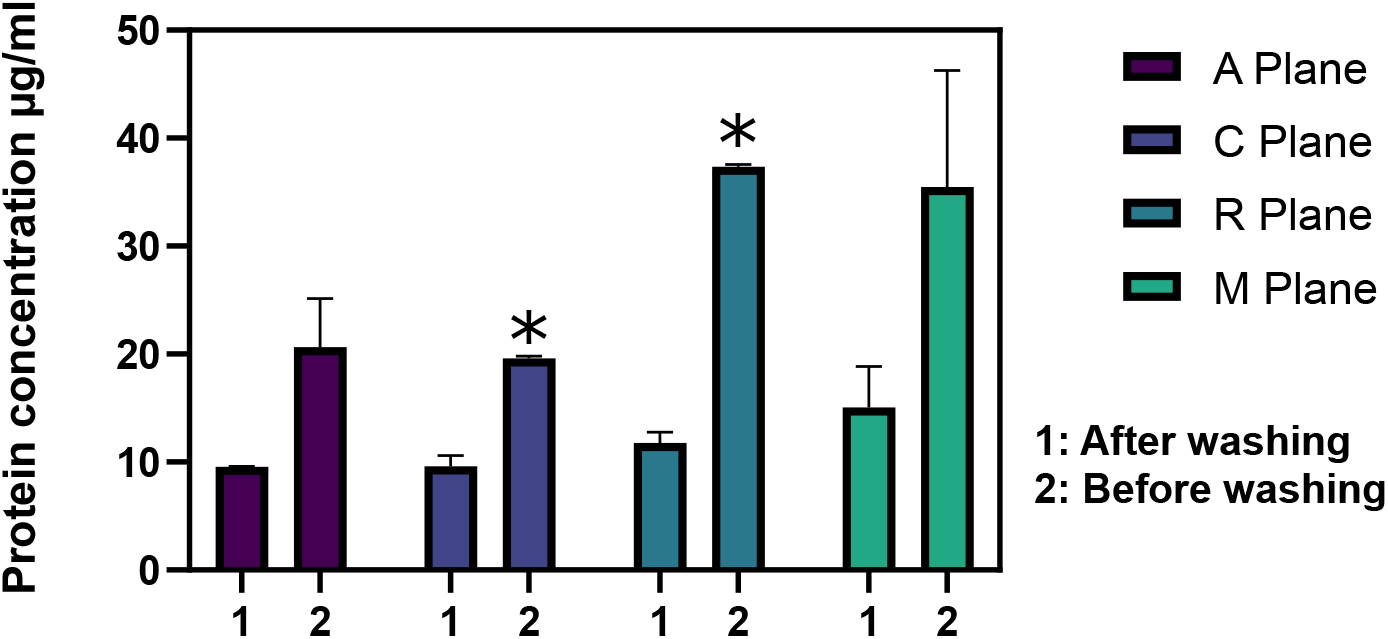
Adsorbed proteins measured by Micro BCA protein adsorption kit before (right) and after washing (left) on alumina planes (n = 3, *p = 0.02).

## Discussion

In this research, we aimed to investigate the influence of the crystallographic orientation on hemocompatibility by examining interactions of blood cells, protein adsorptions, and endothelial cells with different planes of single crystals of alumina having different characteristics, e.g., chemical structure, roughness, electrical potential, surface energy and charge. Our discussion involved two main cellular responses: endothelialization and platelet activation. Additionally, we considered the proteins which are momentarily adsorbed to the material’s surface after the blood contact [28, 29]. Because, it is clear that the initial interactions between the surface and the proteins tailor the platelet and endothelial cell responses. The driving forces for protein adhesion are reported as electrostatic, van der Waals and hydrolyzation interactions [30]. Hence, it would be elucidative to initiate the discussion with the molecular interactions at the surface.

To define the surface interactions, specifically on *α*-alumina it is essential to review its hexagonal lattice. The model of crystal structure and the main crystallographic planes are shown in Figure 1. The distribution of Al^3+^ and O^2−^ ions on the different planes in a 3D model was illustrated. The decisive factor of material properties can be attributed to these atomic arrangements and atomic planar density. When the surface is exposed to the atmosphere, hydrogen bonding develops thin water films on the surface with various coordination. The development of the few layers of water mainly determines the solubilization of ions, hydroxyl groups, and other functional groups on the surface, which in turn, influence the dynamics at the interface [31]. The coordination of hydroxyl groups on different crystallographic planes was previously studied by Barron and Zhu *et al*. [32, 33]. It is proposed that oxygen vacancies have a remarkable importance in promoting active sites for the adhesion of functional groups [33]. Among the investigated planes, the C-plane has the highest number of dangling bonds and atomic density (1.3×10 15 at/cm^2^), followed by the M-plane, (2.6×10^14^ at/cm^2^) and A-plane (1.6×10^14^ at/cm^2^) [34]. Planes having high densities of atom steps, kinks, edges and dangling bonds present higher chemical activity [35]. As experimentally confirmed, C-plane has the highest surface energy with the lowest contact angle (Figure 3), indicating more disrupted intermolecular bonds with higher affinity to attach OH groups on the surface.

Zeta potential measurements showed that the A-plane promotes the adhesion of functional groups on the surface with a different chemical reactivity concerning the other planes. As represented in Figure 5, the curve of the A-plane shows significantly higher IEP than the other curves. The shift of the IEP and the presence of the two plateaus are attributed to the presence of both acidic and basic functional groups with a small prevalence of the basic ones. According to the literature [36–38], these functional groups ascribed to be hydroxyls. The chemical reactivity of a OH group is strictly related to the coordination state of the oxygen ion that is bonded to a single octahedral or tetrahedral Al^3+^ ion, i.e., two or three aluminum ions, respectively. Five different OH groups with different chemical reactivity are identified in the literature on alumina: basic groups (pKa around 9), weakly basic (pKa at 6.6-6.8), almost neutral (pKa at 4.4-4), and acidic (pKa around 3) ones [36, 37]. Multiply coordinated hydroxyls have a low pKa (acidic reactivity), while singly coordinated surface hydroxyl groups have a high pKa (basic reactivity). A prevalence of singly coordinated and basic groups on plane A and of almost neutral OH groups on the other planes can be inferred from the zeta potential titration curves. Furthermore, the absolute value of zeta potential at the physiological pH of plane A was significantly lower than the other planes.

We aimed to gain deeper insight into level of impurities and adsorbed species at the surface by SIMS analysis. Our findings revealed distinct behaviours of different crystallographic planes concerning OH^−^ peaks (Figure 6). 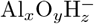 secondary ions are not ‘structure-related clusters’ but rather result from combinations of ionized Al_2_O_3_ components and water molecules, these secondary ions are likely linked to –OH groups [39]. The varying intensities observed are thus attributed to differences in hydration among the sapphire surfaces.

At first glance, the SIMS results appeared to contradict our wettability data, as the C-plane exhibited higher wettability than the M- and R-planes. However, considering surface roughness, this can be explained by the Cassie-Baxter effect, in which rougher surfaces trap air bubbles, effectively reducing the contact angle [40]. It is also important to note the limited number of replicates in SIMS measurements and the uncertainty regarding the crystallographic effects in SIMS results [41]. Nonetheless, the intensity order of 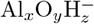 signals across all four orientations was consistent with platelet behaviour (Figure 9).

The density and the variety of functional groups determine the immediate adsorption of plasma proteins onto the biomaterial surface followed by their conformational changes. This phenomenon occurs through various physical interaction forces such as van der Waals forces and hydrogen bonding. Vroman effect discusses the sequence of protein adsorption from high molecular weight to low molecular weight [42]. This effect is primarily observed on the hydrophilic surfaces which is correlated with our SIMS analysis. The relatively high intensity of signals at the higher mass range for the spectra of R-, M-, and C-planes signifies higher adsorption of small-weight organic molecules with putative hydrophobic alkyl groups. The lowest intensities of the hydrophobic impurity peaks on the C-plane were consistent with its high wettability (Figures 3 and 6). Overall, these findings suggest that SIMS analysis serves as a valuable complement to other methods, providing additional insights into surface chemistry. However, our current results are preliminary; further, more comprehensive investigations will be followed.

Similar to the platelets, the acceleration of endothelial cell layer formation on biomaterials primarily occurs through the action of adhesive biomolecules. We observed higher HCAECs proliferation and confluence on the A-plane. These results were further corroborated by pFAK staining which explores mature focal adhesions between HCAECs and their surrounding microenvironment. Qualitative analysis showed adhesive points were spread over a larger area on the A-plane surface with a high density (Figures 7 and 8).

Now we elaborate the discussion with surface potential, which influences the protein adhesion and cell behaviours. Despite to strong charging characteristics of alumina, the KPFM maps on different planes significantly differ indicating higher potential on A plane than the other planes (Figure 4). Promoted HCAEC adhesion on A plane with the highest surface potential correlates the previous studies [30, 43] and verifies that the cell adhesive properties can be dynamically switched by tuning the surface potential. In contrast to HCAECs, platelet adhesion on the surface is inhibited with elevated surface potential which suggests higher hemocompatibility.

Finally, we briefly address the role of surface roughness, which can exert multifunctional effects depending on the rugophilicity of the cell type [44]. Increased surface roughness can promote cell adhesion by providing a larger surface area and favorable topographical features, which may lead to enhanced cellular attachment and higher proliferation rates. On the other hand, smoother surfaces are often associated with improved cell spreading and stable attachment due to reduced steric hindrance and enhanced surface contact. In the present study, the maximum roughness difference observed among the different crystallographic planes (Figure 2) was below nm, a range that was intentionally selected to enable the investigation of other surface parameters independently. When compared with the average size of platelets (∼2 *µ*m), such minor roughness differences are not expected to promote blood cell activation or adhesion. Consistent with this expectation, our results did not reveal a linear relationship between these small variations in surface roughness and cell adhesion.

## Conclusion

In this study, we investigated the impact of crystallographic orientation on cellular behaviours, focusing on the crystal structure of alumina single crystals. Our findings highlighted the complex interplay between various parameters within the system, including protein adsorptions, blood components, and endothelial cells, each exhibiting unique characteristics that dynamically interact with their surroundings. Our results underscored the significant influence of crystallographic planes on thrombogenic response. Low platelet activation and higher endothelial proliferation on specific planes verify that surface orientation is an essential parameter that cannot be overlooked in the in vitro examination of blood-contacting materials. This significant influence is attributed to the differences in the stacking order of surface atoms that modulate adhered functional species, consequently altering surface properties, particularly potential and energy.

## Acknowledgments

The authors express gratitude to Mr. Alejandro Gomez and Mr. Philipp Schraeder for their assistance with the introduction to pFAK analysis, and Mr. Philipp Jacobs for conducting the XRD measurements. The contribution of all blood donors is deeply appreciated. This study received financial support from the Deutsche Forschungsgemeinschaft (DFG, German Research Foundation) - grant number 405895710.

